# Matrix Nanoscale Mechanics Regulates Exosome Production by Mesenchymal Stem Cells

**DOI:** 10.1101/2023.09.19.558518

**Authors:** Alexandra Chrysanthou, Julien E. Gautrot

**Author notes:** Correspondence: Julien E. Gautrot.

## Abstract

Complex biotherapeutics such exosomes offer attractive opportunities for cell-free treatment of disease and conditions difficult to address with single, defined compounds. However their production remains challenging as adherent cells proposed to secrete therapeutic extra cellular vesicles require scalable platforms. In addition, the role of biomaterials design parameters on processes regulating vesicular secretory phenotypes is unclear. Here we propose the use of bioactive microdroplets, or bioemulsions, as microcarriers for the culture of mesenchymal stem cells and production of exosomes. We demonstrate 100% increase in the output of extracellular vesicles on bioemulsions. The impact of matrix mechanics on this process is then investigated, and in particular interfacial shear mechanical properties of corresponding liquid-liquid interfaces forming microdroplets. We find that such local nanoscale mechanics regulates not only cell adhesion, but also exosome output. We find that exosomes generated by cells cultured on bioemulsion microdroplets retain a high content of protein and RNA cargos. Finally, we demonstrate that the cold-shock protein YBox 1, previously associated with RNA packaging, is modulated by matrix mechanics and regulates exosome production. Together, these results demonstrate the impact of local interfacial mechanics on the adhesion and secretory machinery and provide a proof of concept for the application of bioemulsions for the production of complex biotherapeutics.

## Introduction

Stem cell-based therapies may allow tackling a wide range of conditions and diseases, whether based on mesenchymal stromal cells of various origins or whether based on induced pluripotent stem cells [1–3]. Underlying this regenerative potential, the secretory phenotype of these cells, in particular associated with pro-angiogenic factors, has been proposed to play a central role in tissue regeneration [4–6]. Extra-cellular vesicles (EVs) such as exosomes have emerged as important biotherapeutics orchestrating reparative processes and, in turn, present important potential for cell-free stem cell therapies [7]. Indeed, such approach enables harnessing some of the crucial regenerative potential of stem cells whilst minimising risks associated with cell therapies, such as tumorigenicity and immunogenicity.

EVs are classified according to two sub-categories, based on their biogenesis pathways [8]: exosomes range in size from 30 to 150 nm and originate from multi-vesicular bodies (MVBs), while microvesicles are larger (size range of 100-1000 nm) and bud directly from the cell membrane [8,9]. Exosome biogenesis is initiated in the endosome, followed by maturation in the late endosome and MVBs. Accumulation of intraluminal vesicles through endosomal membrane invagination leads to fusion with the cell membrane and release of exosomes in the extracellular space. These processes are orchestrated by a complex molecular machinery, within which ESCRT and heat shock proteins (e.g. Hsp90) play a central role, regulating invagination, fusion, budding and pinching events [8,10]. In addition, GTPases such as Rab proteins regulate transportation of MVBs along the actin and microtubule cytoskeleton during exosome genesis [11,12]. Maturation is also associated with the recruitment of specific proteins and biomacromolecules often used as exosomal markers, such as tetraspanins CD63 and CD9 and micro RNAs [13]. Finally, although molecules such as Y-box protein 1 (Ybox1) have recently been implicated in miRNA sorting within exosomes [10][14–17], detailed mechanisms controlling the packaging of biomacromolecules within exosomes remain unclear.

The control of exosomal production and composition indeed appears essential, given the role of exosomes in distant signalling, for example in pathological scenarios such as metastasis [12,18,19], in which changes in exosome production and matrix stiffness correlate. Indeed, matrix stiffness was found to promote exosome secretion and induce a more aggressive tumour phenotype [20]. This was found to be mediated through an Akt-Rab8 activation mechanism, directly linked to focal adhesion response to an increase in modulus of the matrix. Such processes not only upregulated exosome production, but also impacted their composition, for example their enrichment in Jagged1, in turn impacting signalling through the Notch pathway. Similarly, exosome production was enhanced in breast cancer cells cultured on stiff matrices and resulted in activation of cell motility and matrix degradation, priming for invasiveness [21]. Matrix stiffness and geometry were shown to regulate vesicular trafficking and exocytosis, through RhoA and Akt activation [22,23]. The exosomes secreted by periodontal ligament stem cells in response to mechanical stretch were found to display altered composition (in particular miRNA content), compared to unstretched samples, in turn affecting the phenotype of targeted cells (in this work, MSCs) [24]. Surprisingly, exosome production by MSCs was recently found to be upregulated on soft hydrogels (therefore impacting output), without impacting on exosomal composition and therapeutic potential [25]. However, detailed mechanisms via which MSC exosome production is regulated by matrix mechanics and how this may modulate composition and therapeutic cargo packaging remain unclear. The impact of nanoscale mechanics and mechanical heterogeneity or anisotropy, which are increasingly recognised to play a central role in the regulation of adherent cell phenotype [26], on EV production have also not been investigated.

Therefore, the mechanical properties of the matrix are emerging as important parameters enabling the regulation of exosome and EV output, and should be engineered in the design of platforms enabling the production scale up of these cell-free complex therapeutics. In this respect, 3D bioreactors making use of microcarriers to support cell adhesion (often in the context of mesenchymal stromal cells), cell expansion and exosome release have received significant attention [27–29]. Indeed, an important issue for the translation of exosomes for regenerative medicine remains their large-scale production [7,27]. The low yields associated with exosome production limit the expected biological outcome from such therapies and pre-clinical studies have indicated that effective therapeutic dosages in mice require 10^9^–10^11^ EVs scales (corresponding to 10 – 100 μg of exosomal proteins), while most studies report less than 1 μg/mL protein yields [28,30,31]. As a consequence, such high quantities of EVs require the culture of high cell densities and associated costs of medium and microcarriers (for 3D bioreactor production) [9,32].

Microdroplet technologies constitute an emerging platform for the culture and scale up of adherent cell expansion. Indeed, protein assemblies allowing the stabilisation of microdroplets can be engineered to form crosslinked networks at the liquid-liquid interface, resulting in the formation of protein nanosheets with strong nanoscale (interfacial) mechanical properties [33–36]. When combined with their functionalisation with bioactive moieties, for example peptide or ECM proteins ligating integrin receptors, protein nanosheet-stabilised microdroplets sustain the adhesion, spreading and expansion of adherent cells, including keratinocytes, mesenchymal stem cells, induced pluripotent stem cells and HEK293 cells [37–43]. Mesenchymal stromal cells adhering and proliferating at the surface of protein-nanosheet stabilised liquid interfaces were found to retain the expression of key stem cell markers and their multipotency, even upon long term culture [37]. In addition, these cells retained a secretory phenotype and conditioned a microenvironment that primed the formation of biomimetic adipose-rich bone marrow niches, enabling the expansion of hematopoietic stem cells in vitro and their scale up in conical flask bioreactors [44]. Beyond fluorinated oils routinely used in microdroplet technologies, for example for single cell RNA sequencing [45–47], a broader range of silicone, mineral and plant based oils have also been applied for the culture of MSCs [48], enabling the use of biomedical grade materials for cell expansion. Therefore, bioactive emulsions, or bioemulsions (emulsions based on microdroplets stabilised by surfactants presenting direct bioactive properties), offer advantages over solid or hydrogel microcarriers, which are difficult to process, relatively costly and can introduce microplastic contaminants in the cell products. However, the ability of bioemulsions to support the production of complex therapeutics such as exosomes and EVs, and how the nanoscale mechanics of corresponding protein nanosheet-stabilised interfaces, has not been investigated yet.

In this report, we propose the use of bioemulsions for the production of MSC-derived exosomes and extracellular vesicles. The nanoscale mechanical properties of corresponding liquid-liquid interfaces stabilised by protein nanosheets are characterised and the impact of the strong nanoscale shear mechanics of these interfaces on MSC adhesion and EV production is investigated. The broader impact of matrix mechanics on EV release is then explored in the context of poly(acrylamide) hydrogels, in comparison to tissue culture plastic and bioemulsions. The role of cytoskeleton tension and assembly on EV production is examined. Finally, the impact of matrix compliance on the composition of EVs is characterised and the mechanism of mechanosensing of associated secretory phenotype, including the role of the translational regulator YBox1, is examined.

## Materials and Methods

### Emulsion preparation and functionalisation

To generate emulsions, 1 mL of fluorinated oil (Novec 7500, ACOTA) and 2 mL of protein solution (β-lactoglobulin; >90%, Sigma, from bovine milk, at a concentration of 1 mg/mL in PBS) were added into a glass vial. The vial was vigorously shaken (Vortex, 10 s) to form the emulsion and left for 1h at room temperature. The upper liquid phase was aspirated and replaced with PBS six times. The droplet surface was functionalised as described before [49]. Briefly, sulfo-succinimidyl-4-(N-maleimidomethyl)cyclohexane-l-carboxylate (sulfo-SMCC, Sigma, at a concentration of 2 mg/mL in PBS) were added to the emulsion and kept to react for 1 h at room temperature. The upper liquid phase was aspirated and replaced with PBS 3 times. The peptide solution (CGGRGDSPG, Proteogenix, free of TFA, 1.6 mg/ mL in PBS; 2 mL) was added to the emulsion and incubated for 1 h at room temperature. The upper liquid phase was aspirated and replaced with PBS 3 times followed by washing (3 times) with cell culture medium. The samples were kept in cell culture medium for at least 30 minutes before cell seeding.

### Polyacrylamide (PAAm) hydrogels formation

The PAAm hydrogels were prepared and characterised as described previously [50,51]. Briefly, air plasma-activated coverslips were functionalised with methacrylate residues by incubation in solutions of 3-(trimethoxysilyl)propyl methacrylate (90 μL, Fisher) in anhydrous toluene (30 mL, Fisher) overnight, at room temperature, followed by washing with copious amounts of ethanol and water. Then, 40 % acrylamide (Sigma) and 2 % bis-acrylamide (Sigma) stock solutions were mixed at ratios of 1:250 and 3:160 (acrylamide:bisacrylamide) to form gels of 2 kPa and 115 kPa stiffness’s, respectively. PAAm gels were formed on the functionalised coverslips of 13 and 35 mm diameter in order to be placed in 24– and 6-well plates, respectively. Before cell seeding, the hydrogels were kept in 1 % antibiotic solution (Penicillin-Streptomycin 5,000 U/mL; Thermo Fisher) at 4⁰C overnight. Hydrogels were washed extensively with PBS and activated via photoirradiation (365 nm, 0.8 mW for 5 minutes) with sulfosuccinimidyl-6-4’-azido-2’-nitrophenylamino hexanoate (Sulfo-SANPAH; Thermo Scientific Pierce). To do so, 20 μL of sulfo-SANPAH stock solution (DMSO 50 mg/mL) was diluted into 980 μL deionised water. A volume of 100 μL or 300 μL was added to each gel (volumes corresponding to 24– or 6-well plates, respectively). Hydrogels were washed 6 times with sterile PBS and kept in fibronectin solution (20 μg/ mL; Sigma) for 1 h at room temperature.

### Mesenchymal Stem Cell Culture and Seeding

Mesenchymal stem cells (P3−5, Promocell) were cultured in mesenchymal stem cell growth medium 2 (PromoCell). For proliferation assays, MSCs were harvested with Accutase solution (Promocell, 2.5 mL for a T75 flask) containing 0.5 mM EDTA (PromoCell) and incubated at 37 °C for 5 min. Cells were resuspended in medium at a ratio of 1:1, centrifuged for 5 min at 1200 rpm, counted, and re-suspended in MSC medium at a desired density. Cells were allowed to adhere and proliferate in an incubator (37 °C and 5% CO2) for different periods of time (three, five and seven days of culture), prior to staining and imaging. For the isolation of extracellular vesicles, cells were seeded in a 6-well plate (in total 8 wells to have a total surface area 75 cm^2^). Cells were seeded at a concentration of 250,000 cells/well, so that cells were near 70 % confluent after 24 h, when the medium was replaced with serum-free medium (Thermo Fischer; StemPro™ MSC SFM, A1033201) for 48 h. EVs were isolated at day 3 and the wells were refilled with fresh MSC SFM. The same procedure was repeated at days 5 and 7 of culture. For passaging, cells were re-seeded at a density of 300,000 cells per T75 flask.

### Characterisation of particle sizes and concentrations

To characterise the size and concentration of EVs, nanoparticle tracking analysis (NTA) was carried out using a Nanosight LM20 (Nanosight Technology, Malvern). Triplicates of standard measurements (60 seconds per measurement) video captures were carried out (camera level 16, detection threshold 5 for all samples). The data were analysed using the NTA software version 3.2. EVs size were also characterised by dynamic light scattering (DLS). Three samples were measured (in triplicates each) per condition, using a Malvern Zetasizer Nano ZS.

### EV Isolation

For the isolation of EVs, the cell culture medium was changed to serum-free medium for 48 h, when cells were approximately 70 % confluent. The medium was collected from the samples and cleared from cell debris with multiple centrifugation steps at 4⁰C (300 x g for 10 minutes followed by 2,000 x g for 20 minutes and 10,000 x g for 30 minutes)[52]. Supernatants were collected, aliquoted and stored at – 80⁰C. EVs were isolated using the ExoCap Streptavidin Kit (Caltag MedSystems Ltd, MEX-SA), following the protocol recommended by the supplier. Briefly, streptavidin magnetic beads were first conjugated with the desired biotinylated antibody (anti-CD63, biotin; Miltenyi Biotec) and allowed to incubate with the sample stirring overnight, at 4⁰C. Samples were washed 3 times with the washing buffer and the beads were re-suspended in the appropriate buffer for further analysis.

### EV content characterisation

EVs were isolated on streptavidin magnetic beads as described above and the bicinchoninic acid (BCA) Protein Assay Kit (Thermo Fischer, 23225) was used to characterise their protein content. The magnetic beads with captured EVs were washed once with cold PBS, and then lysed in RIPA buffer (Thermo Fischer, 89900) on ice for 10 minutes. The samples were mixed several times and kept at the magnetic rack for 1 minute. Supernatant were collected and centrifuged for 15 min at 14,000 x g. Supernatants were discarded and the pellet was re-suspended in PBS. BCA solution were added to the samples and kept at 37⁰C for 3 h. The adsorption of the samples were measured at 562 nm with a spectrophotometer.

### Treatment with disruptors of cytoskeleton assembly

Cell were seeded on the substrate of interest, in 24-well plates (at density of 30,000 cells per well). The inhibitors Y-27632 (Sigma) and Jasplakinolide (Sigma) were diluted in DMSO at a concentration of 10 mM. Inhibitors solutions were diluted in MSC medium (at a concentration of 10 μM) and added to the samples for 4 or 24 h. Samples were then washed once with PBS and fixed for imaging. For the EVs isolation, cells were cultured in 6-well plates (at a density of 300,000 cells per well). After the addition of inhibitor solution, the medium was collected and cleared as described earlier before further analysis.

### Immuno-fluorescence staining

Samples were washed once with PBS and fixed with 4% paraformaldehyde for 10 min at room temperature. Thereafter, samples were washed three times with PBS and permeabilized with 0.2% Triton X100 (Sigma-Aldrich) for 5 min at room temperature. After washing with PBS (three times), samples were blocked for 1 h in 3 % bovine serum albumin (BSA). The blocking buffer was partly removed from the emulsion samples; not allowing them to be exposed to air, and all samples were incubated with primary antibodies at 4°C overnight. Samples were washed six times with PBS and incubated for 1 h with the secondary antibodies in blocking buffer (BSA 3 % in PBS; phalloidin, 1:500; DAPI, 1:1000; vinculin, 1:1000; CD63, 1:500; CD9, 1:500). After washing with PBS (six times), emulsion samples were transferred to Ibidi wells and gels on coverslips were mounted on glass slides for imaging. The same procedure was followed for the captured EVs on magnetic beads, using the magnetic rack for the washing steps. SYTO RNA Select Green Fluorescent Cell Stain (Thermo Fischer) was added to the samples after imaging with the secondary antibodies. An amount of 100 μL of labelling solution (500 nM staining solution) was added to each sample and incubated for 20 minutes at room temperature.

### Immuno-Fluorescence Microscopy and Data Analysis

Confocal microscopy images were acquired with a Zeiss 710 confocal microscope. To determine the cell spreading areas, images (phalloidin staining of the actin cytoskeleton) were analyzed by thresholding and watershedding in Fiji ImageJ. A total number of 50 cells per conditions were analysed at day one and three of culture.

### Transfection assay

MSCs were cultured in 6-well plates (300,000 per well) and in 24-well plates on coverslips (100,000 per well) and allowed to adhere for 24 h. Transfections were carried out with siRNA (Thermo Fischer, Silencer Select siRNA) at a concentration of 30 pmol/ well. siRNA and Lipofectamine RNAiMAX (Thermofischer, 13778150) were diluted separately in OPTI-MEM (Thermo Fischer, 31985062). The diluted siRNA and the diluted Lipofectamine were then mixed gently in 1:1 ratio and incubated for 20 minutes at room temperature. The medium from each well was aspirated and washed with PBS and replaced with 400 μL OPTI-MEM. 100 μL of the siRNA-lipid complex were added drop by drop to the cells and mixed gently. The plates were incubated at 37°C for 4 h. The medium was aspirated and each well was washed once with PBS before they were refill with MSC medium. The coverslips in the 24-well plates were fixed and stained for imaging after 24 h. The imaging was carried out with a Leica DMI8 and the transfection efficiency was quantified using Fiji ImageJ. Silencing sequences and controls are as follows, from Thermo Fisher: Silencer™ Select Negative Control (4390843) and Silencer® Select siRNAs for YBX1 (siRNA IDs s9732 and s9733) and YBX2 (siRNA IDs s27393 and s27394).

### Statistical Analysis

Statistical analysis was carried out using OriginPro 9 through one-way ANOVA with Tukey test for posthoc analysis. Significance was determined by * P < 0.05, ** P < 0.01, *** P < 0.001 and n.s., non-significant. A full summary of statistical analysis is provided in the supplementary information.

## Results and Discussion

To support the growth of MSCs on scalable bioemulsions, microdroplets stabilised by β-lactoglobulin (βLG) were investigated (Figure 1A) [49]. As this protein is not bioactive, at least from the point of view of integrin ligation and the promotion of cell adhesion, the resulting droplets were further functionalised with RGD peptides. Protein nanosheets are required to accomplish a triple role for bioemulsion design, conferring stability owing to their tensioactive properties, cell adhesion through integrin-binding peptides and scaffolding properties, in order to resist cell-mediated forces. It was found that, even in the absence of additional co-surfactants, which had been found critical to regulate the viscoelastic properties of albumin nanosheets, βLG formed comparatively elastic interfaces [36]. This was confirmed by interfacial rheology (Figure 1B), indicating a rapid increase in interfacial shear storage and loss moduli upon injection of βLG. The two moduli rapidly diverged, with the storage component dominating near 8-fold over the loss component. After applying a protocol allowing the exchange of the aqueous phase to remove free protein from the solution, the addition of sulfo-SMCC led to a further 3-fold increase in interfacial shear modulus.

**Figure 1.**
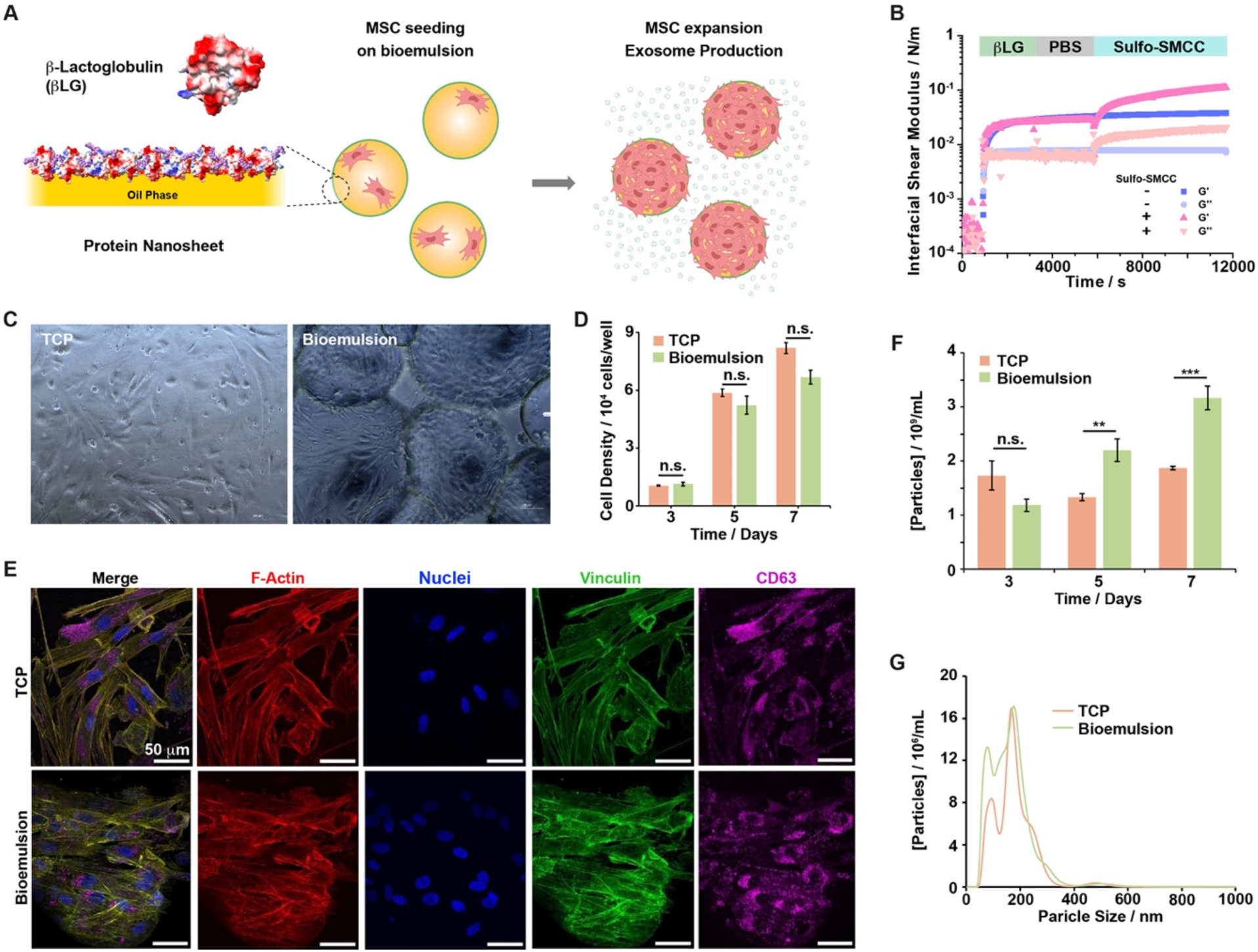
A) Schematic representation of bioemulsions formed microdroplets stabilised by β-Lactoglobulin nanosheets, allowing the expansion of MSCs and promoting the production of exosomes. B)Formation and mechanical properties of β-Lactoglobulin nanosheets at oil-water interfaces (Novec 7500/PBS), with and without treatment with sulfo-SMCC (2 mg/ mL). Time sweeps of the evolution of interfacial shear storage and loss moduli following injection of β-Lactoglobulin (1 mg/mL final concentration) at 900 s time point. Oscillating amplitude of 10^−4^ rad and frequency of 1 Hz. C) Images of MSCs after seven days of growth on tissue culture plastic (TCP) and β-lactoglobulin nanosheet-stabilised bioemulsion. D) Densities of MSCs proliferating on TCP and bioemulsion, quantified at different time points. E) Confocal microscopy images of MSCs grown on TCP and bioemulsion after seven days of culture (blue, DAPI; red, phalloidin; green, vinculin; magenta, CD63). F) Concentration of EVs secreted by MSCs cultured on TCP and bioemulsion at different time points. G) Corresponding size profiles at day three of culture. Error bars are s.e.m.; n = 3.

These results were further confirmed by frequency sweeps and stress relaxation experiments, which confirmed the formation of stiff elastic interfaces following sulfo-SMCC treatment (Supplementary Figure S1-2). This behaviour suggests that sulfo-SMCC, upon coupling through its activated succinimidyl ester residue, enables further reaction and crosslinking via maleimides. This may be attributed to remaining free cysteines on the βLG proteins, or could be associated with reactivity with remaining lysines and other nucleophiles, via Michael addition. Alternatively, the addition of the relatively hydrophobic cyclohexyl groups may be sufficient to induce physical crosslinking. Beyond enabling bioconjugation with cell adhesive peptides, sulfo-SMCC may therefore also result in the crosslinking of protein nanosheets and strengthening of associated interfacial mechanics. With an ultimate interfacial shear modulus near 100 mN/m, and assuming a thickness near 10 nm [36], the equivalent bulk Young’s modulus of βLG-SMCC interfaces is estimated to be near 30 MPa, significantly above the minimum range of moduli of hydrogels supporting MSC adhesion [50,53].

MSCs were next seeded on the resulting βLG-RGD bioemulsions and cultured for 7 days (Figures 1A and C). The proliferation of these cells was monitored during this time period and compared to that of MSCs adhering to tissue culture plastic (TCP, Figure 1D). MSCs expanded significantly during this time period, both on TCP and βLG-RGD bioemulsions, with expansion rates slightly higher in the former case, although this was not statistically significant. In agreement with the impact of cell adhesion and cytoskeleton assembly on MSC fate and cycling [50,54,55], cells adhering to both TCP and βLG-RGD bioemulsions displayed clear focal adhesions and stress fibres, with a well-structured actin cytoskeleton (Figure 1E).

In order to evaluate the output of extra-cellular vesicles (EVs) as a function of culture substrate, cells were cultured in cleared medium (free from debris and vesicles or particulates contained in serum), prior to monitoring of particle densities, associated with EV secretion (Figure 1F). A significant increase in EV secretion and accumulation into the culture medium was observed in the case of MSCs cultured on βLG-RGD bioemulsions (2 fold compared to TCP). In addition, capture by magnetic microparticles and immunostaining confirmed that EVs released were positive for the exosome markers CD63 and CD9 (Supplementary Figure S3). When corrected for differences in cell densities, this corresponds to a 100 % increase, compared to TCP. This increase was associated with increased densities of CD63+ vesicles, in bioemulsion cultures (Figure 1E), without significant change in the size distributions (Figure 1G). Therefore these results suggest that substrate nanoscale mechanics not only impacts on cell adhesion, spreading and proliferation at liquid-liquid interfaces, but also extracellular vesicle production and associated signalling.

The significant increase in EV production in bioemulsion cultures is in agreement with growing evidence that a number of physical microenvironment parameters regulate both vesicular membrane activity and extra-cellular vesicle production [20,22,23]. The observation that the production of EVs by MSCs was increased at liquid-liquid interfaces raised the possibility that matrix stiffness promoted enhanced EV output by MSCs.

To examine the role of matrix compliance on vesicle production, MSCs were cultured on soft (2 kPa) and stiff (115 kPa) poly(acrylamide) (PAAm) hydrogels functionalised with fibronectin (Figure 2). Cell adhesion and spreading were significantly impacted by hydrogel stiffness, in agreement with well-established literature [50,53]. 24 h after seeding, whereas cells adhering to 115 kPa PAAm hydrogels displayed a well-structured actin cytoskeleton and mature focal adhesions, MSCs adhering to 2 kPa hydrogels did not (Figure 2A). In addition, although CD63+ vesicles could be clearly observed both on soft and stiff hydrogels, immunostainings microscopy indicated relatively aggregated structures in cells adhering to 2 kPa PAAm gels, whereas they were more homogenously distributed in the case of cells adhering to 115 kPa gels. This observation is likely associated with the significant change in cytoskeleton organisation observed in response to matrix compliance, and associated change in membrane tension, vesicular transport and exocytosis [22,23].

**Figure 2.**
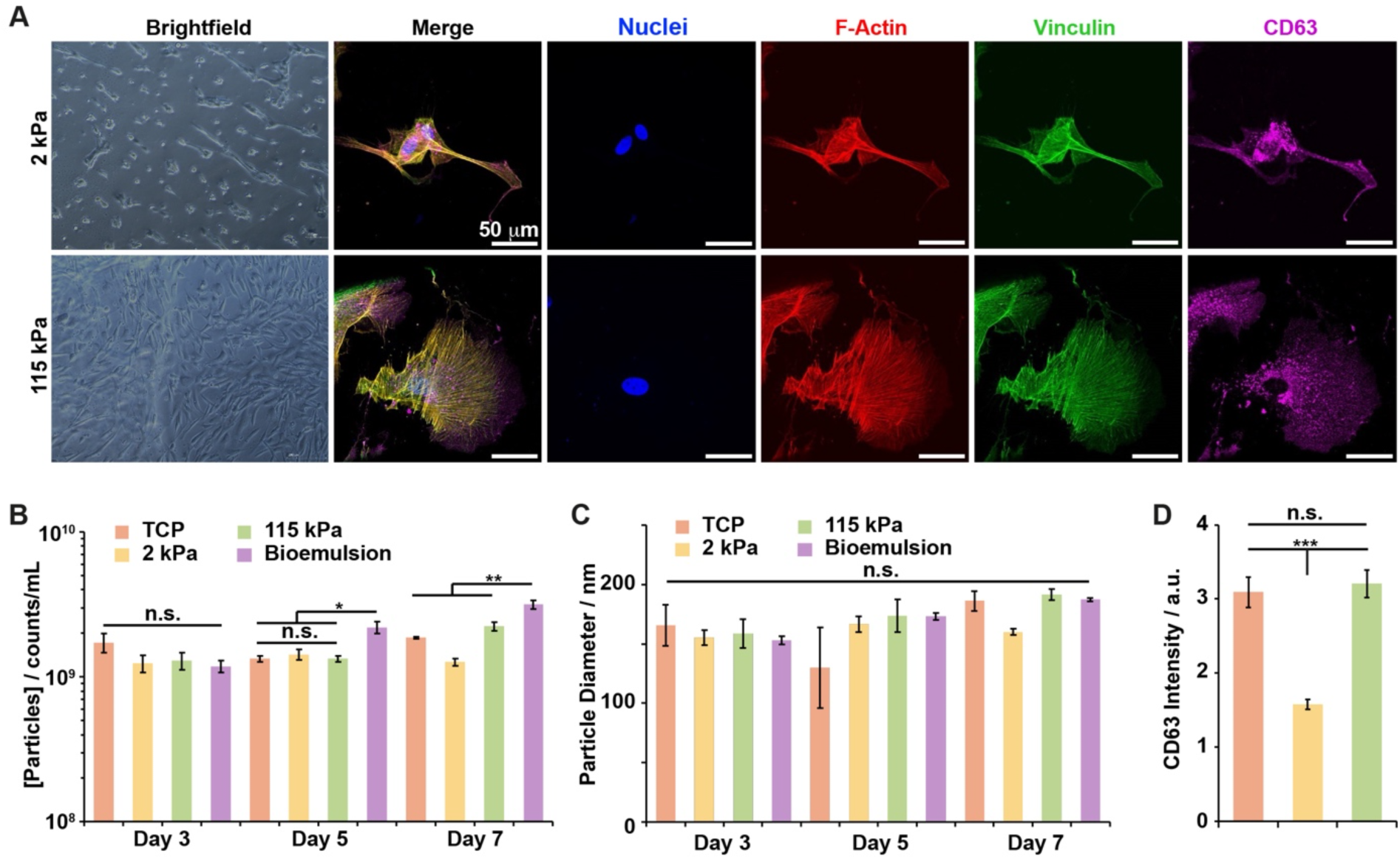
Substrate stiffness impacts cell morphology and EVs secretion. A) Bright field and confocal microscopy images of MSCs cultured on the soft (2 kPa) and stiff (115 kPa) PAAm hydrogels (bright field, day 7 of culture; confocal microscopy, 24 h). B) Concentration of EVs secreted from TCP, PAAm hydrogels (2 and 115 kPa) and bioemulsion on day three, five and seven of culture. C) Corresponding sizes of secreted EVs. D) Intensity of CD63 + objects (total intensity per cell) presented by cells cultured on TCP, soft and stiff hydrogels (2 and 115 kPa, respectively) after seven days of culture. Scale bars are 50 μm. Error bars are s.e.m.; n = 3.

The impact of matrix stiffness on EV production was next investigated, over 7 days of culture. At early time points (day 3), despite the restricted cell spreading observed on softer hydrogels, the concentrations of EVs released were found to be equivalent on all substrates, although slightly higher on TCP (Figure 2B). However, by day 7, more significant differences could be seen, with MSCs cultured on 2 kPa PAAm gels displaying significantly fewer EVs compared to cells growing on 115 kPa PAAm gels and TCP. Interestingly, cells growing on bioemulsions displayed higher EV production compared to cells cultured on stiff hydrogels. The sizes of the vesicles secreted were found to be comparable in all conditions, in the range of 130-200 nm (Figure 2C). At day 7, EV production also correlated with the expression and presentation of CD63 + particles, based on fluorescence images (Figure 2D), confirming the particle counts determined. Therefore, the strong nanoscale mechanical environment associated with protein nanosheets assembled at liquid-liquid interfaces was found to dominate cell response to the mechanics of their micro-environment and sustained not only cell spreading, but also promoted enhanced EV production.

Considering the role played by cytoskeleton assembly and contractility on the regulation of MSC phenotype, migration, cytokine expression and exocytosis, the impact that such processes could have on EV production was next examined. After initial adhesion to TCP and PAAm hydrogels with 2 and 115 kPa Young’s moduli, MSCs were treated with the Rho kinase (ROCK) inhibitor Y27632 and the inhibitor of actin disassembly Jasplakinolide [56], followed by 24 h of further culture, immunostaining and imaging (Figure 3A). In line with previous observations, such treatment had a significant impact on cell adhesion and cytoskeleton assembly. In both cases, cell spreading was found to be independent from matrix stiffness, with apparent cytoskeletal disruption (Figure 3A and Supplementary Figure S4).

**Figure 3.**
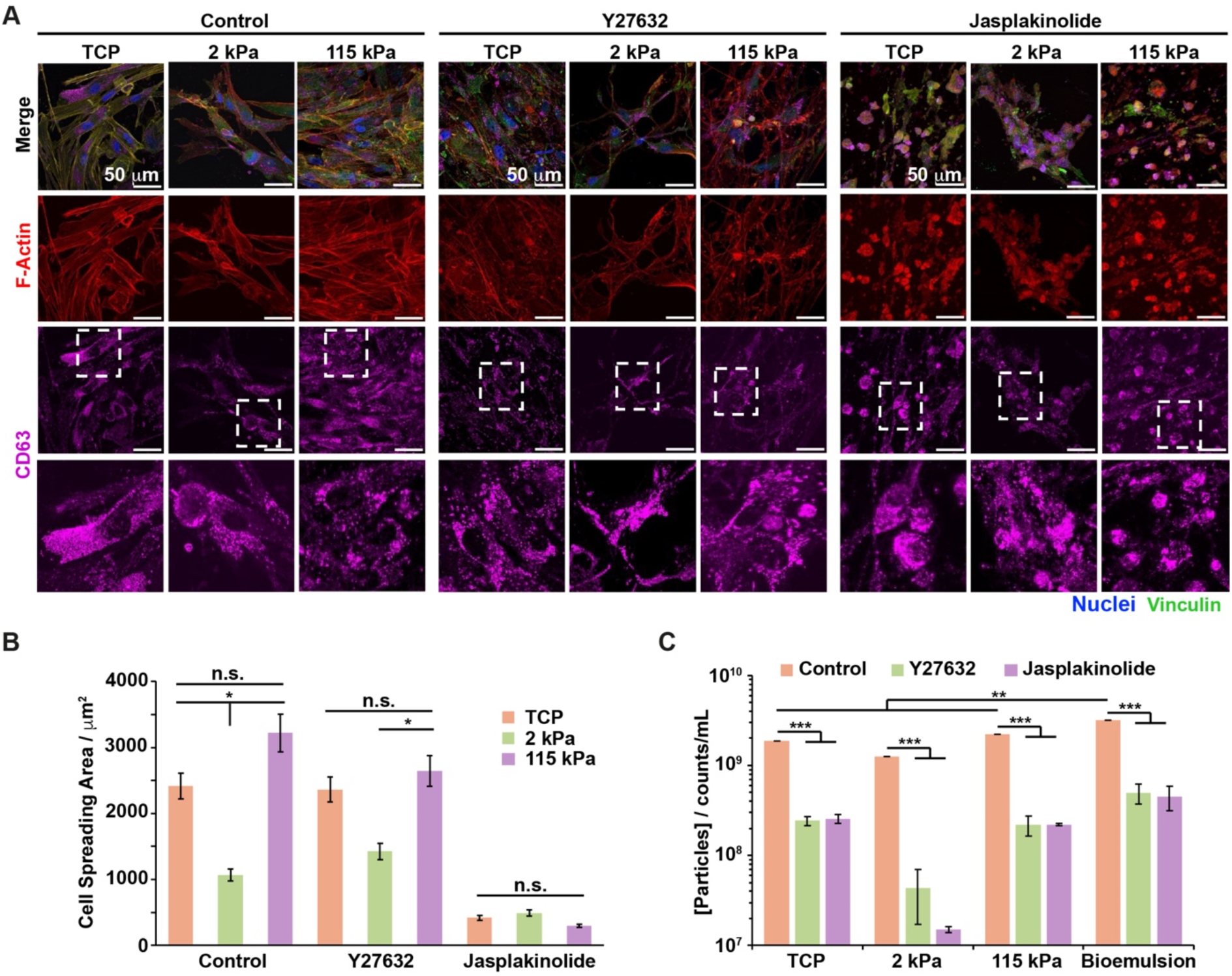
Actin cytoskeleton disruption impacts on the secretion of EVs. A) Confocal microscopy images of MSCs cultured on TCP and PAAm hydrogels (2 and 115 kPa), treated with the ROCK inhibitor Y27632 and Jasplakinolide (blue, DAPI; red, phalloidin; green, vinculin; magenta, CD63). B) Changes in cell spreading as a result of Y27632 and Jasplakinolide treatment, after 24 h of culture. C) Concentration of EVs secreted by MSCs cultured on TCP, PAAm hydrogels (2 and 115 kPa) and bioemulsions, after 24 h of Y27632 and Jasplakinolide treatment. Scale bars are 50 μm. Error bars are s.e.m.; n = 3.

However, whereas MSCs treated with Y27632 displayed high spreading on all substrates, cell spreading was found to be significantly restricted by Jasplakinolide treatment. Stress fibres were disrupted in cells treated with Y27632, but large actin clusters were observed for cells treated with Jasplakinolide. In all cases, CD63+ structures could be clearly observed in cells treated with these compounds, irrespective of the substrate on which they were cultured (Figure 3A). The localisation of associated putative CD63+ vesicles was however significantly affected by disruption of the cytoskeleton organisation, with images implying aggregation and clustering, likely reflecting significant perturbation of vesicle transport. In turn, EV production was significantly impacted by actin disruption (Figure 3C). In all cases, EV concentrations measured upon culture in medium supplemented with actin disruptors were nearly 1 order of magnitude lower than upon culture on the same substrate, in the normal culture medium. This reduction was particularly striking in the case of MSCs treated with Jasplakinolide when cultured on 2 kPa PAAm gels. Therefore, these results are consistent with an important impact of cytoskeleton assembly on EV production, likely through combined effects of such disruption on vesicle transport and membrane tension, two processes regulating exocytosis [12,22].

The impact of substrate mechanics on extravesicular protein and RNA content was quantified next. After recovery from different culture substrates, EVs were lysed and their total protein content analysed. Although protein contents were found to be relatively equivalent at early time points, differences were more apparent at days 5 and 7, with protein concentrations overall significantly higher on bioemulsions compared to those harvested from rigid TCP or compliant PAAm hydrogels (Figure 4A). Similarly, RNA content (quantified via fluorescence microscopy directly capture of EVs by magnetic microparticles) was also found to be enhanced on bioemulsions (Figures 4B and C). In addition, RNA content was reduced more significantly on softer (2 kPa) PAAm hydrogels. However, when corrected for differences in EV concentrations (Supplementary Figure S5), these differences did not persist and the estimated protein concentration per EV was found to be comparable, indicating that EV content remained globally unchanged, although total output was affected by matrix stiffness.

**Figure 4.**
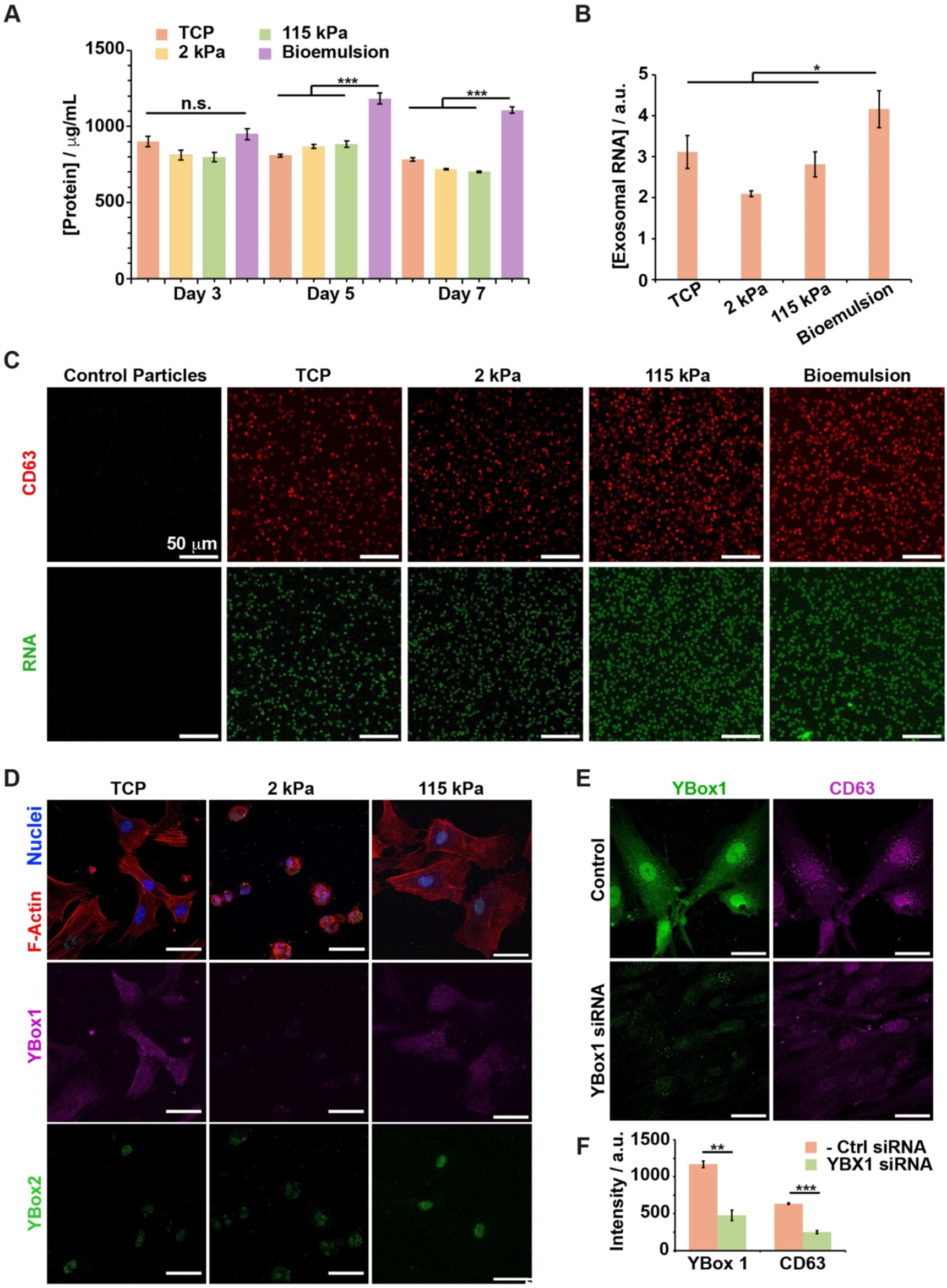
Substrate nanoscale mechanics impacts EVs composition. A) Protein content of EVs isolated EVs from MSCs cultured on TCP, PAAm hydrogels (2 and 115 kPa) and bioemulsions at day three, five and seven. B) Quantification of exosomal RNA recovered from corresponding substrates at day 7. C) Confocal microscopy images of labelled exosomal RNA from EVs (captured on magnetic beads; red, CD63; green, RNA) harvested from on TCP, PAAm hydrogels (2 and 115 kPa) and bioemulsions. D) Fluorescence microscopy images of YBox 1 and YBox 2 expression by MSCs cultured on TCP and PAAm hydrogels (2 and 115 kPa). E) Confocal microscopy images confirming the knock down of YBox 1 and its impact on the formation of CD63 + vesicles. F) Corresponding quantification of YBox 1 and CD63 intensities. Scale bars are 50 μm. Error bars are s.e.m.; n = 3.

As micro RNAs are predominant RNAs found in EVs and are packaged within exosomes through interactions with the translational regulator YBox1 [14,16,17], the impact of matrix stiffness on YBox1-mediated RNA packaging was next examined. To investigate a possible involvement of YBox1 in the matrix-mediated modulation of EV production, immunostaining of YBox1 and YBox2, another cold-shock protein implicated in the regulation of RNA translation [57], but not previously associated with RNA packaging, was examined (Figure 4D). YBox1 was found to be predominantly cytoplasmic, with some nuclear overlap, but presumably reflecting peri-nuclear association, rather than nuclear localisation. In contrast, YBox2 was found to be predominantly nuclear or peri-nuclear. In response to matrix compliance, the pool of YBox1 was significantly depleted in MSCs adhering to 2 kPa PAAm hydrogels, compared to 115 kPa PAAm gels or TCP. In the case of YBox2, expression levels were also found to be significantly reduced on 2 kPa PAAm hydrogels compared to 115 kPa PAAm or TCP. Therefore the expression level and localisation of YBox proteins 1 and 2 were found to be regulated by matrix compliance, suggesting a potential role of these proteins in the regulation of exocytosis, EV production and RNA packaging by matrix mechanical properties.

In order to investigate the impact of YBox proteins 1 and 2 on EV production, their knock down was implemented, prior to examination of CD63 expression and localisation via immuno-staining (Figure 4E and Supplementary Figures S6 and S7). The expression of both proteins was reduced by 80-90% using two independent siRNA targets (see methods for details). However, whereas this led to a clear reduction in CD63+ staining intensity in YBox1 knock down, compared to non-targeting controls (Figure 4E), YBox2 knock down had a more modest impact on CD63 expression or apparent localisation (Supplementary Figure S6). Therefore, these data suggest that, in addition to the regulation of exocytosis via membrane tension, matrix compliance may also regulate YBox1 expression and localisation and in turn EV production and RNA packaging.

## Conclusions

Exosomes and EVs are increasingly considered as a viable cell-free strategy for regenerative medicine, but require development of manufacturing pipelines in order to cope with the throughput associated with the safe translation of these technologies. The significant enhancement of EV production on bioemulsions may contribute to bring these complex therapeutics closer to the clinic, in a scalable, easily processable and safe format. Beyond the selection of oils, proteins and co-surfactants that may be applied for the design of bioemulsion platforms, the understanding of the impact of nanoscale mechanics on MSC secretory phenotype, EV release and packaging of associated biotherapeutics will be essential to optimise the performance of these systems. Combined with state-of-the-art strategies for cell engineering and expression of defined biotherapeutics, bioemulsions may help to bring precise therapeutics design and production closer to translation.

This report also further demonstrate the important role that local, nanoscale, mechanics plays for the regulation of cell phenotype. Despite the low viscosity of the oil used in this study, the nanoscale mechanics of protein nanosheet interfaces was found to support cell spreading, proliferation and to enhance EV production. Hence, whereas PAAm hydrogel compliance significantly reduced EV outputs, protein nanosheet-stabilised microdroplets promoted higher levels of secretion compared to relatively stiff hydrogels and rigid TCP. In this respect, the role of nanoscale mechanics on the regulation of exocytosis and RNA/protein packaging within EVs remains unclear. However, this report proposes that YBox1 expression and sub-cellular localisation is regulated by matrix compliance and controls the packaging of EVs and potentially RNA cargos. Although the precise molecular mechanism via which such process could be regulated remains to be established, the potential synergy with membrane tension-mediated exocytic control may underpin overall EV production and therapeutic packaging. We could not observe any correlation between the actin cytoskeleton and YBox1 localisation, despite reports suggesting potential association of related P50 with actin[58]. Proteins regulating focal adhesion or cytoskeleton assembly, such as zyxin which was reported to associate with Ybox1 to modulate downstream translation of RXRγ mRNA [59], could also regulate EV packaging. Elucidating the detail of these mechanisms could prove particularly important to further design biomaterials for EV production, as well as to understand how microenvironmental factors may mediate pathological scenarios through EV signalling.

## Acknowledgement

We thank Dr Ioanna Keklikoglou for insightful discussions of our work and manuscript. We thank the European Research Council (ProLiCell, 772462) for support.

