## Supplementary Information for "Matrix Nanoscale Mechanics Regulates Exosome Production by Mesenchymal Stem Cells"

Alexandra Chrysanthou[1] and Julien E. Gautrot[1*]

[1] School of Engineering and Materials Science, Queen Mary University of London, Mile End Road, London E1 4NS, United Kingdom.

* Correspondence:

Julien E. Gautrot

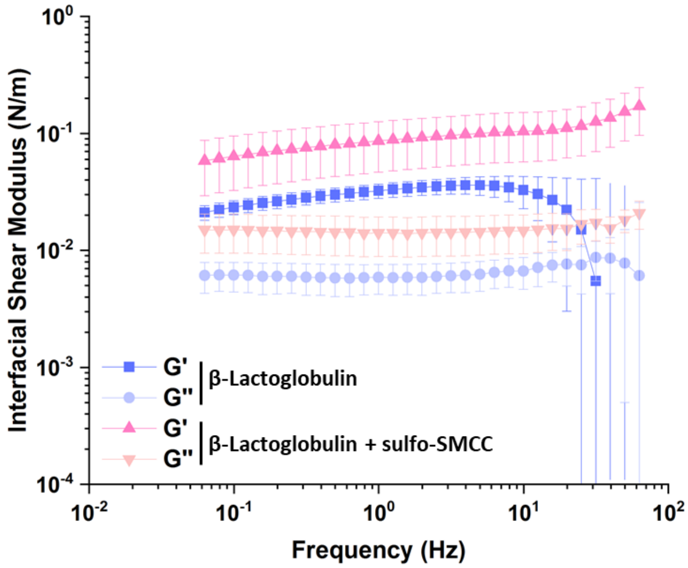


**Supplementary Figure S1.** Frequency sweep at oscillating amplitude of $10^{-4}$ rad carried out for the characterization of β–Lactoglobulin (all at 1 mg/mL) with and sulfo-SMCC (at 2mg/ mL). All experiments were carried out at interfaces between PBS and Novec 7500 (fluorinated) oil. Oscillating amplitude of $10^{-4}$ rad.

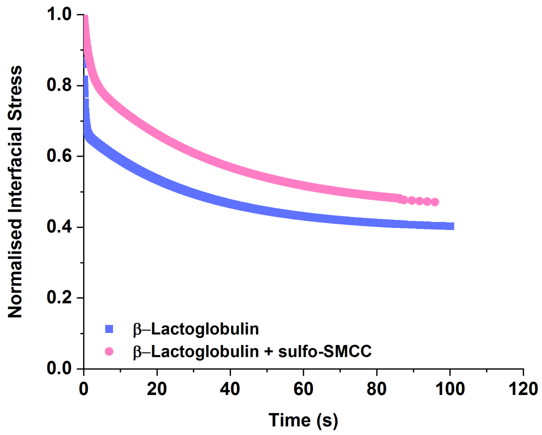


**Supplementary Figure S2.** Stress relaxation experiments carried out at 0.5 % strain on protein nanosheets stabilised with β-Lactoglobulin at liquid-liquid interfaces with and without sulfo-SMCC (2 mg/mL).

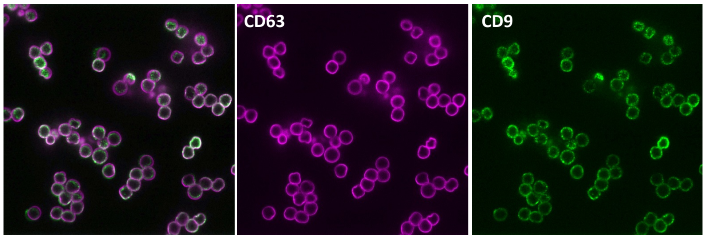


**Supplementary Figure S3.** Representative confocal microscopy images of EVs captured on magnetic beads (magenta, CD63; green, CD9).

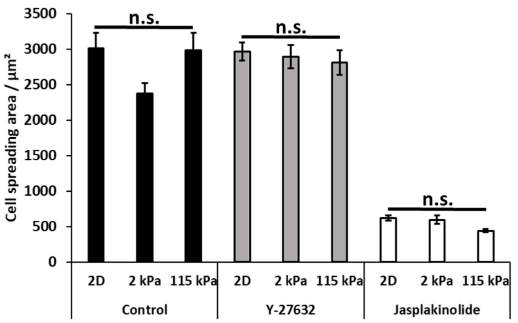


**Supplementary Figure S4.** Changes in cell spreading as a result of Y27632 and Jasplakinolide treatment, after 3 days of culture.

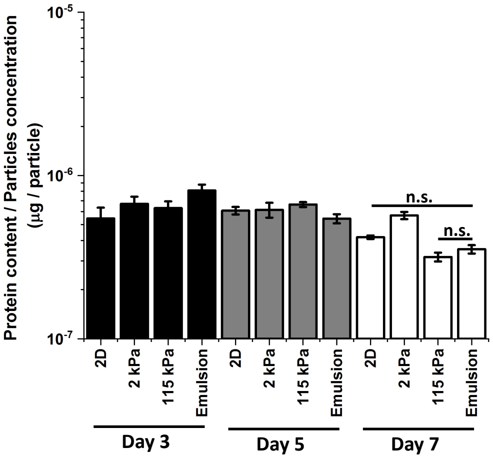


**Supplementary Figure S5.** Protein concentrations of EVs secreted by MSCs (relative to the particle concentration) cultured on a TCP, PAAm hydrogels (2 and 115 kPa) and bioemulsions after three, five and seven days of culture. Error bars are s.e.m.; n = 3.

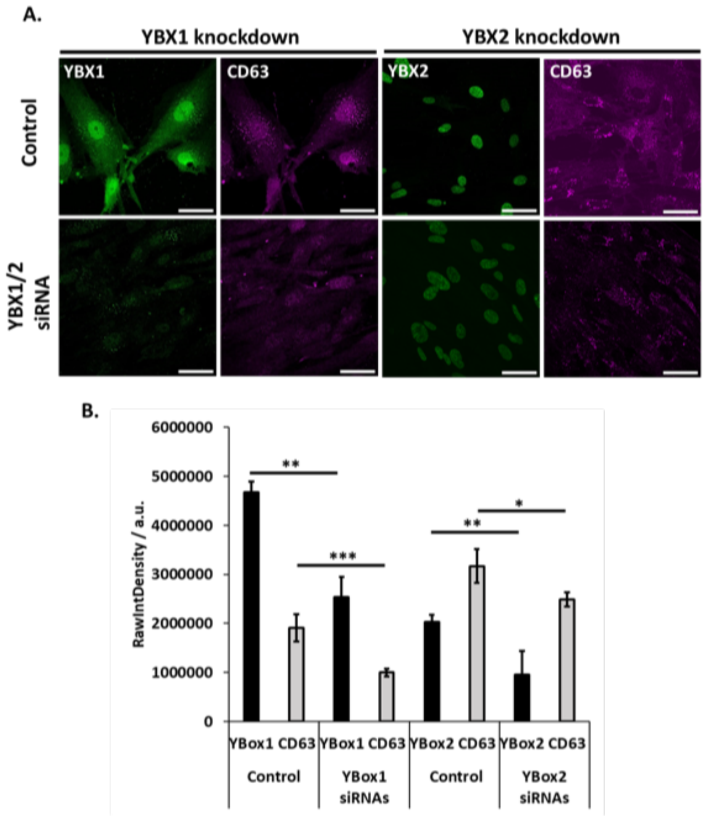


**Supplementary Figure S6.** A) Confocal microscopy images of Ybox 1, Ybox 2 and CD63 immuno-stained samples in which cells were treated with corresponding siRNA (negative controls vs YBX1 or YBX2) for 24 h (magenta, CD63; green, YBX1, YBX2). B) Corresponding quantification of the raw integrated density (per cell) before and after the knockdown for YBX1 and YBX2. Scale bars are 50 µm. Error bars are s.e.m.; n = 3.

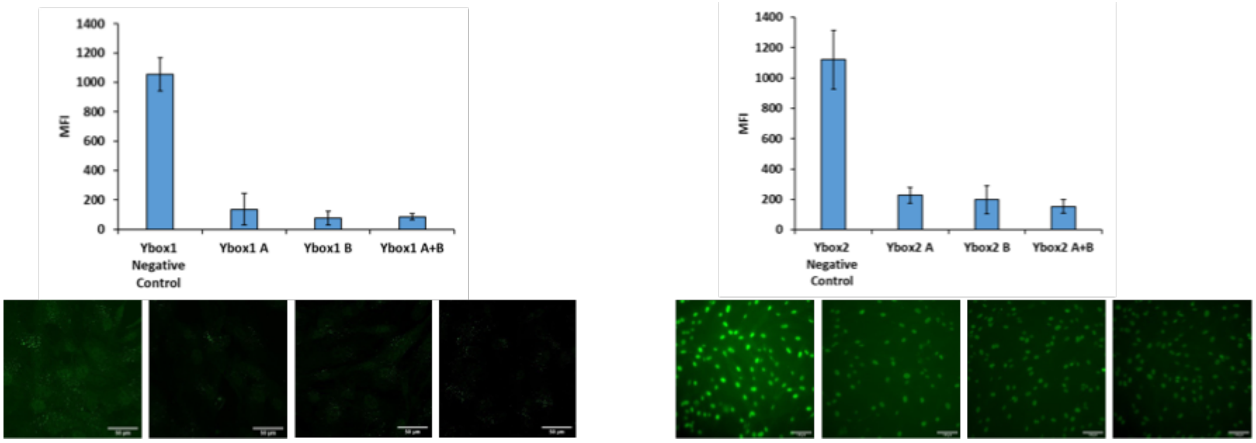


**Supplementary Figure S7.** Quantification of Ybox 1 and Ybox 2 expression in response to corresponding siRNA knockdown. Corresponding mean fluorescence intensities. Scale bars are 50 µm. Error bars are s.e.m.; n = 3.

**Supplementary Table S1.** Summary of statistical significance of the interfacial shear storage moduli obtained from frequency sweeps (frequency of 1 Hz).

| | MeanDiff | Prob | |
|---|---|---|---|
| b-LG+sulfoSMCC b-LG | 0.10449 | 0.01592 | * |

**Supplementary Table S2.** Summary of statistical significance of the interfacial shear loss moduli obtained from frequency sweeps (frequency of 1 Hz).

| | MeanDiff | Prob | |
|---|---|---|---|
| b-LG+sulfoSMCC b-LG | 0.01142 | 0.02119 | * |

**Supplementary Table S3.** Summary of statistical analysis of protein concentrations of EVs secreted by MSCs cultured on TCP, PAAm hydrogels (2 and 115 kPa) and bioemulsions at day three. Protein content was estimated after correction with particle concentrations at the same time point.

| | MeanDiff | Prob | |
|---|---|---|---|
| 2kpa TCP | -395370 | 0.6027 | n.s. |
| 115kpa TCP | -300821 | 0.77037 | n.s. |
| 115kpa 2kpa | 94549.45 | 0.98946 | n.s. |
| Emulsion TCP | -674127 | 0.2109 | n.s. |
| Emulsion 2kpa | -278757 | 0.80657 | n.s. |
| Emulsion 115kpa | -373306 | 0.64248 | n.s. |

**Supplementary Table S4.** Summary of statistical analysis of protein concentrations of EVs secreted by MSCs cultured on TCP, PAAm hydrogels (2 and 115 kPa) and bioemulsions at day five. Protein content was estimated after correction with particle concentrations at the same time point.

| | MeanDiff | Prob | |
|---|---|---|---|
| 2kpa TCP | 3005.176 | 1 | n.s. |
| 115kpa TCP | -137919 | 0.86173 | n.s. |
| 115kpa 2kpa | -140924 | 0.85426 | n.s. |
| Emulsion TCP | 210789.7 | 0.64858 | n.s. |
| Emulsion 2kpa | 207784.5 | 0.6581 | n.s. |
| Emulsion 115kpa | 348708.5 | 0.27426 | n.s. |

**Supplementary Table S5.** Summary of statistical analysis of protein concentrations of EVs secreted by MSCs cultured on TCP, PAAm hydrogels (2 and 115 kPa) and bioemulsions at day seven. Protein content was estimated after correction with particle concentrations at the same time point.

| | MeanDiff | Prob | |
|---|---|---|---|
| 2kpa TCP | -622782 | 0.07659 | n.s. |
| 115kpa TCP | 792277.3 | 0.02538 | * |
| 115kpa 2kpa | 1415059 | 7.77E-04 | *** |
| Emulsion TCP | 474576.5 | 0.20007 | n.s. |
| Emulsion 2kpa | 1097358 | 0.00405 | ** |
| Emulsion 115kpa | -317701 | 0.4907 | n.s. |

**Supplementary Table S6.** Summary of statistical analysis of data obtained by epifluorescence images for YBox1 knockdown efficiency and the impact on CD63.

| | MeanDiff | Prob | |
|---|---|---|---|
| Ybx1after Ybx1before | -2765329.167 | 0.00104 | ** |
| CD63before CD63after | 1539997.667 | 9.19E-05 | *** |

**Supplementary Table S7.** Summary of statistical analysis of data obtained by epifluorescence images for YBox2 knockdown efficiency and the impact on CD63.

| | MeanDiff | Prob | |
|---|---|---|---|
| Ybx2after Ybx2before | -1077751 | 0.00194 | ** |
| CD63before CD63after | -684114 | 0.02564 | * |

**Supplementary Table S8.** Summary of statistical analysis of data quantifying protein concentrations of EVs secreted by MSCs cultured on TCP, PAAm hydrogels (2 and 115 kPa) and bioemulsions at day three. The protein content was estimated after correction with particle concentrations at the same time point.

| | MeanDiff | Prob | |
|---|---|---|---|
| Bioemulsion TCP | 2.64E-07 | 0.08038 | n.s. |
| 2 kPa TCP | 1.24E-07 | 0.33741 | n.s. |
| 115 kPa TCP | 8.51E-08 | 0.47881 | n.s. |
| 115 kPa 2 kPa | -3.91E-08 | 0.70211 | n.s. |
| Bioemulsion 2 kPa | 1.40E-07 | 0.23441 | n.s. |
| Bioemulsion 115 kPa | 1.79E-07 | 0.13052 | n.s. |

**Supplementary Table S9.** Summary of statistical analysis of data quantifying protein concentrations of EVs secreted by MSCs cultured on TCP, PAAm hydrogels (2 and 115 kPa) and bioemulsions at day five. The protein content was estimated after correction with particle concentrations at the same time point.

| | MeanDiff | Prob | |
|---|---|---|---|
| 2 kPa TCP | 6.43E-09 | 0.99949 | n.s. |
| 115 kPa TCP | 5.43E-08 | 0.79483 | n.s. |
| 115 kPa 2 kPa | 4.78E-08 | 0.84725 | n.s. |
| Bioemulsion TCP | -6.53E-08 | 0.69488 | n.s. |
| Bioemulsion 2 kPa | -7.17E-08 | 0.63392 | n.s. |
| Bioemulsion 115 kPa | -1.20E-07 | 0.2542 | n.s. |

**Supplementary Table S10.** Summary of statistical analysis of data quantifying protein concentrations of EVs secreted by MSCs cultured on TCP, PAAm hydrogels (2 and 115 kPa) and bioemulsions at day seven. The protein content was estimated after correction with particle concentrations at the same time point.

| | MeanDiff | Prob | |
|---|---|---|---|
| 2 kPa TCP | 1.51E-07 | 0.62054 | n.s. |
| 115 kPa TCP | -1.02E-07 | 0.94574 | n.s. |
| 115 kPa 2 kPa | -2.53E-07 | 0.05759 | n.s. |
| Bioemulsion TCP | -6.53E-08 | 0.99824 | n.s. |
| Bioemulsion 2 kPa | -2.16E-07 | 0.16135 | n.s. |
| Bioemulsion 115 kPa | 3.68E-08 | 0.99999 | n.s. |
